# A Spatiotemporal Proteomic Map of Human Adipogenesis

**DOI:** 10.1101/2023.07.01.547321

**Authors:** Felix Klingelhuber, Scott Frendo-Cumbo, Muhmmad Omar-Hmeadi, Austin J. Taylor, Pamela Kakimoto, Morgane Couchet, Sara Ribicic, Martin Wabitsch, Ana C. Messias, Arcangela Iuso, Timo D. Müller, Mikael Rydén, Niklas Mejhert, Natalie Krahmer

## Abstract

White adipocytes function as the major energy reservoir in humans by storing substantial amounts of triglycerides and their dysfunction is associated with metabolic disorders. However, the mechanisms underlying cellular specialization during adipogenesis remain unknown. Here, using a high-sensitivity-high throughput workflow, we generated a spatiotemporal proteomic atlas of human adipogenesis encompassing information for ~8.000 proteins. Our systematic approach gives insights into cellular remodeling and the spatial reorganization of metabolic pathways to optimize cells for lipid accumulation and highlights the coordinated regulation of protein localization and abundance during adipogenesis. More specifically, we identified a compartment-specific regulation of protein levels and localization changes of metabolic enzymes to reprogram branched chain amino acid and one-carbon metabolism to provide building blocks and reduction equivalents for lipid synthesis. Additionally, we identified C19orf12 as a differentiation induced and adipocyte-specific lipid droplet (LD) protein, which interacts with the translocase of the outer membrane (TOM) complex of LD associated mitochondria and regulates adipocyte lipid storage. Overall, our study provides a comprehensive resource for understanding human adipogenesis and for future discoveries in the field.

**KEY POINTS:**

- Human adipogenesis induces distinct temporal changes in protein abundance
- 20% of all detected proteins change organellar localization during adipogenesis
- BCAA and one-carbon metabolism enzyme levels and localizations are coordinately regulated to promote adipogenesis
- C19orf12 is an adipocyte specific LD protein and regulator of lipid storage

## INTRODUCTION

Living organisms have evolved the capacity to store energy in the form of fat in lipid droplets (LDs). The core of these storage organelles contain neutral lipids, such as triglycerides and an average healthy adult stores 10-25 kg of fatprimarily in white adipose tissue (WAT), with each kg equivalent to 9,000 kcal^1^.

WAT characterized by few, but large adipocytes (hypertrophy) is associated with insulin resistance and the secretion of pro-inflammatory cytokines, whereas WAT displaying a higher number of small adipocytes (hyperplasia) is linked to a metabolically healthy phenotype^8–10^. The fact that the balance between hyperplasia and hypertrophy associates strongly with the risk of developing metabolic complications of obesity, underscores the importance to understand how adipocytes acquire their remarkable capacity for lipid storage and mobilization during differentiation, and to identify cellular processes that underly healthy adipogenesis and lipid dynamics.

Recent technological advances in transcriptomics have significantly improved our understanding of adipocyte heterogeneity and transcriptional networks underlying adipogenesis^11–15^. However, it has become increasingly clear that post-transcriptional processes are also critical for regulating protein levels and activity during adipogenesis^16–19^. These dynamic processes, along with the remodeling of organelles can be more thoroughly understood at the proteome level, which remains poorly characterized in the context of adipogenesis. To date, proteomic analyses have provided static snapshots of the protein landscape, but not yet covered the spatiotemporal aspects of adipogenesis^20–23^. As a result, our current understanding of proteomic remodeling and the subcellular organization during adipocyte differentiation remains incomplete.

To address this gap, we have generated a comprehensive spatiotemporal proteomic map across four different human adipocyte models. Our deep and quantitative temporal profiling of protein level changes allowed us to determine proteomic evolution across adipogenesis, which we then compared to primary human white adipocytes and WAT to identify conserved proteomic changes in human adipogenesis. Using a machine-learning-based organelle proteomics approach, we mapped multiple changes in protein localization during adipogenesis. We revealed the coordinated remodeling of metabolic pathways at the level of protein abundance and localization to support de novo lipogenesis, and identified a yet unknown LD protein, C19orf12, as a novel regulator of adipocyte function. We found that C19orf12 localizes to LD-mitochondrial contact regions, and its depletion leads to impaired lipolysis, and increased lipid droplet accumulation. In a human patient cohort, we find *C19orf12* expression in WAT inversely correlated with obesity associated clinical parameters, underlying the key function of C19orf12 in human adipocyte lipid storage. Overall, our study offers a comprehensive resource for understanding the temporally resolved core proteome changes in human adipogenesis, as well as the reorganization of organelles and metabolic pathways that drive human adipogenesis.

## RESULTS

### The temporally resolved core proteome of human adipogenesis

To define the core proteome trajectory during human adipogenesis, we performed liquid chromatography-mass spectrometry (LC-MS) proteomics over the time course of differentiation across different human adipocyte models (Fig. 1A). All models are derived from human adipocyte precursor cells (hAPCs) isolated from the stromal-vascular fraction (SVF) of WAT and have the capacity to differentiate into adipocytes upon treatment with pro-adipogenic cocktails. Two cell types were non-immortalized (SGBS^2^ and hAPC^3,4^), whereas one was immortalized (hWA^5^). To control for the potential effects of the immortalization process, we generated an additional cell model in which hAPCs were immortalized by introducing telomerase reverse transcriptase into the *AAVS1* safe harbor locus using CRISPR/Cas9 engineering (herein termed ThAPC). Thus, a total of four human model systems were included in this study to map the conserved trajectory of adipogenesis independent of cell type-specific effects. Information regarding the origins, immortalization procedures, and differentiation protocols is summarized in Fig.S1A and Table1. For comparison, we also included paired primary samples of subcutaneous abdominal mature adipocytes (pACs), SVF (which contains immature adipocyte precursors), and intact WAT from seven donors (see Methods, Table1). These served as reference points for in vivo adipogenesis.

**Figure 1.**
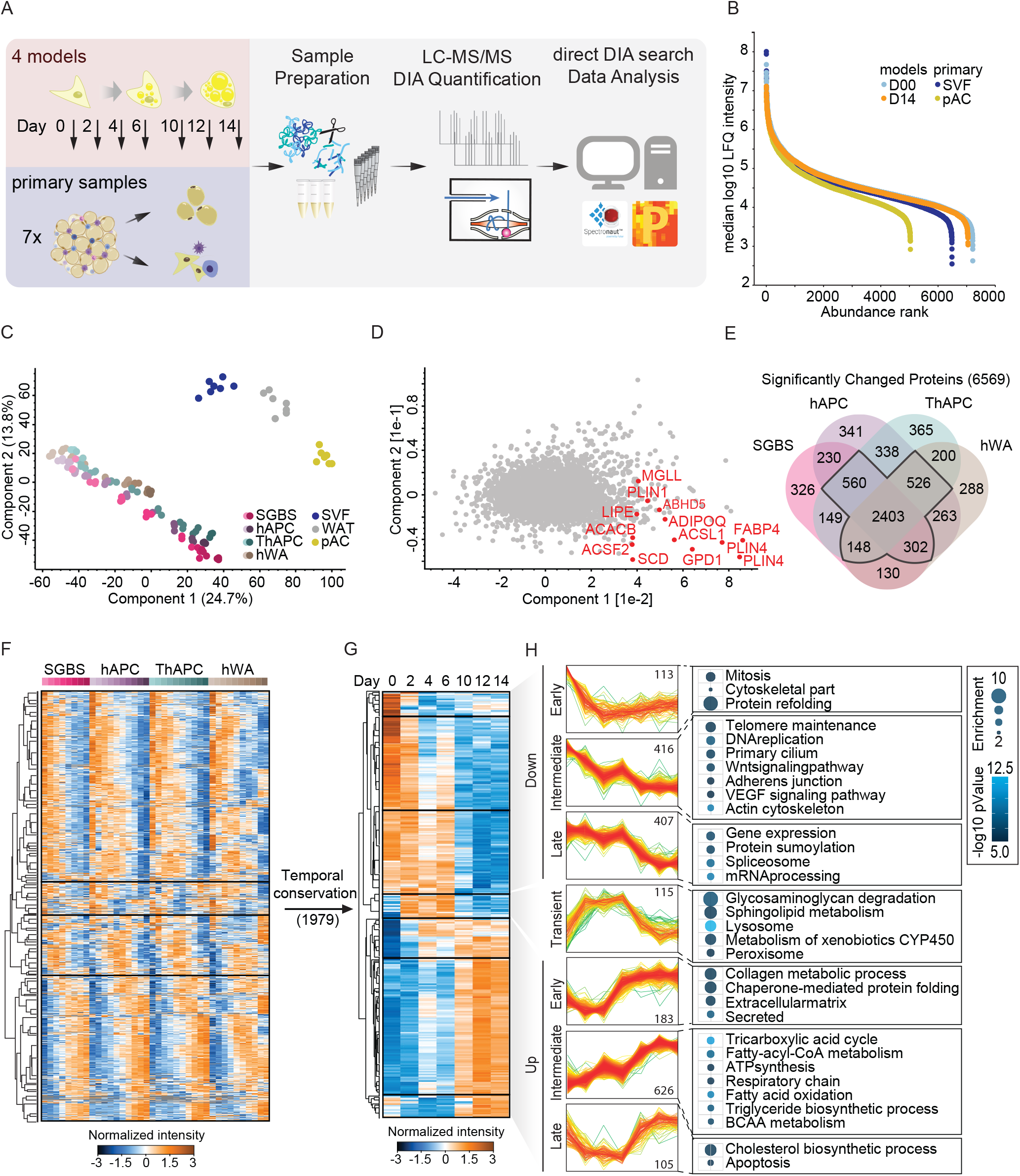
Mapping the temporally resolved core proteome of human adipogenesis. (A) LC-MS based proteomics workflow to map the core proteome of human adipogenesis. Proteomic signatures of 4 human adipogenesis models at seven time points (n=3) were compared with the proteomes from human white adipose tissue and respective pACs and SVFs from 7 patients. (B) Dynamic range of cell line and primary cell proteomes. (C) PCA of primary samples and the adipogenic differentiation stages (depicted as light to dark) of the cell lines. (D) PCA loadings with major driver proteins involved in lipid metabolism shown in red. (E) Overlap of proteins significantly changed over the differentiation in each of the models (individual ANOVA tests, FDR<0.01). (F) Hierarchical clustering of the z-scored temporal profiles of the 3,934 significantly changed proteins in at least three of the four models as outlined in (E). (G) Median temporal profiles across cell models of a subset of (F) with conserved temporal profiles (Pearson Correlation in all comparisons >0). (H) Profiles of individual clusters and selection of enriched annotations (complete list in TableS1). Enrichment values and p values depicted as bubble size and color code, respectively.

**Table 1.** Summary of human adipogenesis models used in the study. Information about donor, origin and initial publication

| Model | Sex | Isolation site | Donor | Immortalized | References |
| --- | --- | --- | --- | --- | --- |
| **SGBS** | m | sc | 4-months old,<br>SGBS syndrome | no | Wabitsch et al.,<br>2001 |
| **hAPC** | m | sc | 16-y, healthy BMI:<br>24 | no | Ehrlund et al., 2013 |
| **ThAPC** | m | sc | 16-y, healthy BMI:<br>24 | TERT | Generated in this<br>study |
| **hWA** | f | sc | 48-y, BMI: 19.9 | TERT | Markussen et al.<br>2017 |

First, we assessed the adipogenic capacity of all four models. By measuring lipid accumulation by BODIPY staining followed by fluorescence microscopy, and mRNA levels of well-established adipogenic marker genes, we found all models to display high differentiation efficiencies (Fig.S1B and Fig.S1C). For proteomic analyses, we sampled the undifferentiated state and at least six time points after induction of differentiation in three biological replicates for each model. Tryptic peptides were analyzed in 1h single shots in data-independent acquisition (DIA) mode (Materials and Methods, Fig.1A). This approach enables higher identification rates over a larger dynamic range and fewer missing values compared to data-dependent acquisition (DDA). Spectronaut analysis quantified 5,979-7,061 protein groups in the four cell models, 3,638-6,403 in the primary samples (pAC/SVF/WAT), and 8,268 in the complete dataset (Fig.S1D). The MS signals spanned an abundance range of five orders of magnitude (Fig.1B) and were highly reproducible among replicates, with an average Pearson’s correlation coefficient of 0.97. We found that a considerable proportion (56-65%) of the proteome underwent remodeling during the differentiation process (ANOVA, FDR 0.01) in all four models. About 6-14% of the quantified proteins showed a more than 10-fold change compared to the undifferentiated state or were exclusive to either the undifferentiated or mature state (Fig.S1E). In total, 3,934 proteins were significantly regulated in at least three of the four models (Fig.1E, F), of which 1,979 proteins displayed a conserved temporal trajectory during adipogenesis (see Methods and Fig.1G). When comparing the samples, the Pearson correlation coefficients were high within the same time points between the systems (Fig.S1F), indicating that there are universal proteomic features of adipogenesis that can be recapitulated in multiple in vitro models. This enabled us to identify the universal events of human adipogenesis, which are not dependent on donors and are not affected by immortalization. Principal component analysis (PCA) showed the temporal trajectory of differentiation states in vitro (the four cell models) and in vivo (SVF vs. pACs) in components 1 and 2, where the adipogenic process of the cell lines projected towards mature adipocytes (Fig.1C). An increase in LD and lipid biosynthesis protein levels was a key factor driving the shift in component 1 (Fig.1D) and we observed a significant increase in protein levels of canonical adipogenesis markers in all four models during differentiation (Fig.S1G). However, our PCA analysis showed that hWA cells reached a less mature state than the other models. This finding is consistent with the reduced expression of classical adipocyte marker genes (Fig.S1C) and lower Pearson correlation coefficients between the proteomes of fully differentiated cell models and pACs (Fig.S1H).

We next performed hierarchical clustering analysis on the conserved temporal profiles. Our results identified distinct clusters that represented early, intermediate, and late responses during adipogenesis (Fig.1H). The early phase was characterized by the downregulation of proteins involved in cell cycle progression, cytoskeletal and extracellular matrix remodeling, secreted factors, and the chaperone system. The transient cluster of temporarily upregulated proteins was characterized by glycosaminoglycan degradation and lysosomal pathways. In addition, the intermediate phase was also defined by a significant increase in enzymes involved in fatty acid and triglyceride metabolism as well as in many mitochondria-related functions that maintain the increased energy demands for triglyceride synthesis, such as the TCA cycle, respiratory chain, and ATP synthesis. Simultaneously, the levels of proteins involved in DNA replication, telomere maintenance, ciliary proteins, and WNT signaling (a pathway that inhibits adipogenesis^6–8^) decreased. In the late phase of adipogenesis, we observed downregulation of spliceosomal and mRNA-processing proteins, and upregulation of cholesterol biosynthesis proteins. Overall, these findings suggest comprehensive remodeling of cellular processes during adipogenesis, with distinct temporal regulation of specific functional pathways.

### A spatial map of adipogenesis

To add a spatial dimension to our proteome of adipogenesis, we utilized protein correlation profiling (PCP), a technique that allows for the analysis of organellar protein localization based on relative abundance profiles. Briefly, for PCP, cells are mechanically lysed, and the organelles are separated by density-gradient centrifugation. Next, proteins are quantified across gradient fractions by LC-MS, and the generated abundance profiles, which are characteristic of the residual cellular compartments, are used to predict protein localization by machine learning^9^.

We applied PCP to mature adipocytes and pre-adipocytes using the SGBS model to identify proteins that displayed different locations during adipogenesis (Fig.2A). 1h LC-MS DIA single shot analyses led to 3,500-5,600 quantified proteins per fraction (Fig.S2A) resulting in cellular maps with higher proteomic coverage and identification rates, less MS runtime, and increased reproducibility compared with traditional DDA-based approaches^10^. A comparison of the median profiles of marker proteins from different cellular compartments indicated clear separation of organelles in our gradients (Fig.2B). Supervised hierarchical clustering of the median profiles from the biological replicates showed distinct clusters for cellular compartments (Fig.2C). By employing support vector machine (SVM)-based supervised learning, we were able to predict primary and potential secondary protein localization using the generated abundance profiles, allowing for the comprehensive annotation of protein distribution. Organellar cluster boundaries were determined using established marker proteins selected based on GO annotations (Methods). Our maps of adipocytes and preadipocytes provide localization information for a total of 7,619 and 8,250 proteins, respectively. Out of these, 4,918 proteins in adipocytes and 4,635 proteins in preadipocytes were confidently assigned to specific organelle clusters using SVMs (see Methods and Fig.2D and 2E). A mean prediction accuracy of 91% was achieved for marker proteins. More than half of the proteins were classified as derived from at least two organelle distributions (Fig.S2B and S2C), consistent with previous studies^11^.

**Figure 2.**
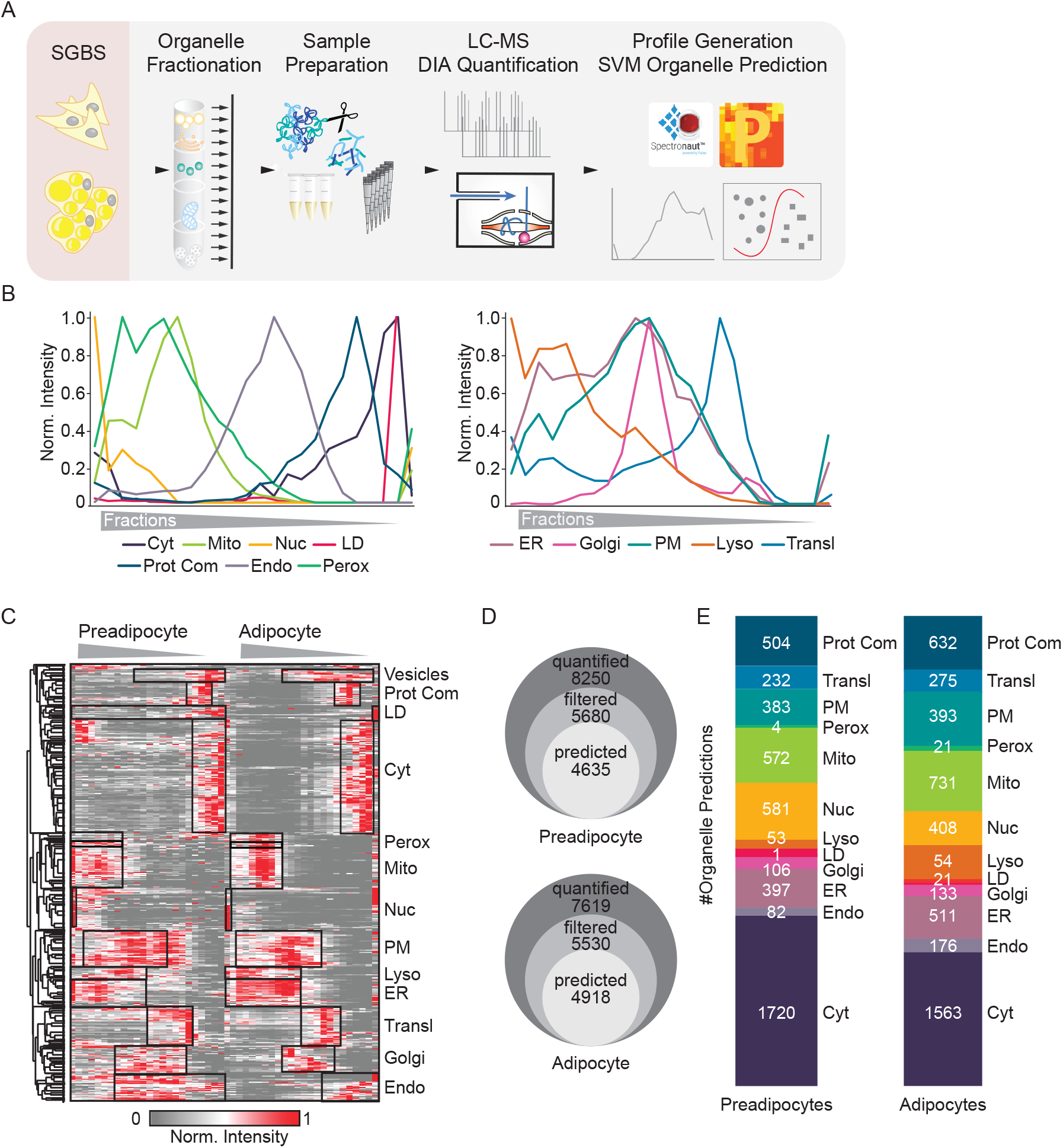
Generation of a cellular map of human adipogenesis. (A) Generation of a human adipocyte organellar map by PCP. Either undifferentiated or fully differentiated SGBS cells were fractionated. Organelle fractions were analyzed by DIA-LC-MS. Protein profiles were generated and SVM-machine learning used to predict protein localizations. (B) Median profiles from biological triplicates for indicated organelles in mature adipocytes based on all proteins assigned to an organelle with a single localization. (C) Supervised hierarchical clustering of protein profiles (median of triplicates) from preadipocytes and adipocytes filtered for inter-replicate correlations >0. GO-terms for organelles enriched in the marked clusters are highlighted. (D) Numbers of quantified, filtered and predicted proteins in preadipocytes and in adipocytes. (E) Numbers of proteins assigned to organelles as first association by SVM-based learning on concatenated protein profiles from quadruplicates.

### Protein localization changes in adipogenesis

SVMs assigned a total of 4,426 proteins with high confidence in both the undifferentiated progenitor cells and the adipocytes, and 898 of them exhibited divergent organelle assignments (Fig.3A). We leveraged these spatial proteomics data and the time-resolved core proteome of adipogenesis to comprehensively characterize organelle remodeling during adipogenesis. By integrating information from both datasets, we were able to predict the proportion of each organelle in the total proteome. Our findings showed that, during adipogenesis, there was an increase in the percentage of mitochondrial, endoplasmic reticulum, endosomal, and LD proteins, whereas the proportion of cytosolic, nuclear, and ribosomal proteins decreased. These changes in organelle composition reflected an overall increase in the total protein mass of all compartments involved in lipid metabolism and secretory functions, which ultimately led to a state that closely resembled the proportional organelle distribution in pACs (Fig.3B, S3A).

**Figure 3.**
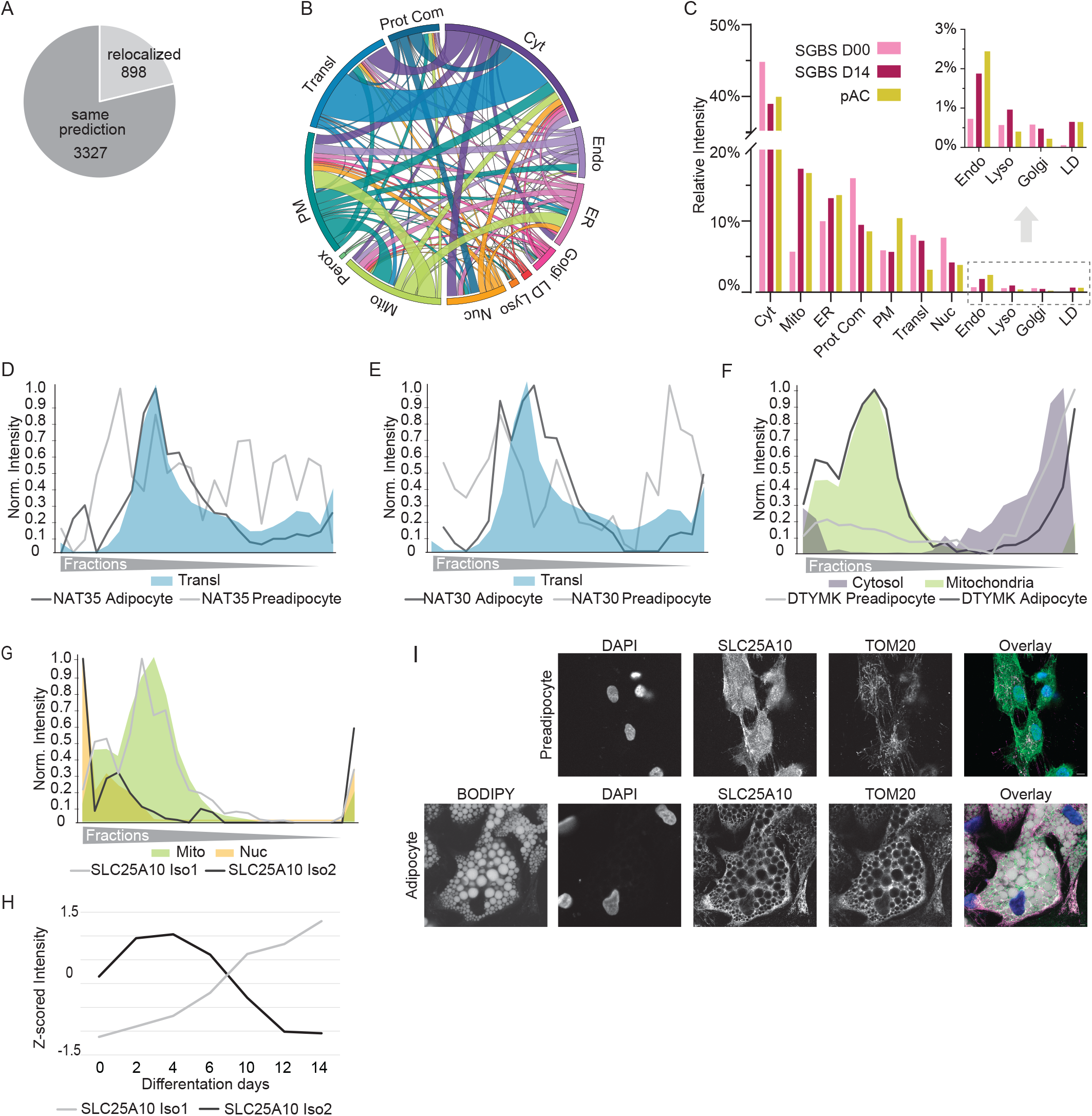
A characterization of protein relocalization events during adipogenesis. (A) Number of proteins assigned to same or different compartments in preadipocytes and adipocytes. (B) Circular river plot displaying relocalization events. (C) Percentage of organelles of total proteome of preadipocytes (SGBS D00), mature adipocytes (SGBS D14) and pACs based on integration of first organelle assignments and summed protein-LFQ intensities in the total proteome analysis. (D) and (E) Profiles of subunits of the NATC complex in preadipocytes and adipocytes overlaid with respective organelle marker profiles. (F) Profile of DTYMK in preadipocytes and adipocytes overlaid with respective organelle marker profiles. (G) Profiles of the two detected isoforms of SLC25A10 in adipocytes overlaid with mitochondrial and nuclear marker profiles. (H) Z-scored median profile of SLC25A10 isoforms from 4 adipogenesis models showing protein levels during adipogenesis. (I) Immunofluorescence for SLC25A10 in preadipocytes and adipocytes, respectively. In the overlay, DAPI is shown in blue, BODIPY in gray, SLC25A10 in green and TOM20 in magenta. Scale bar = 25µm.

Our analysis revealed that certain compartments exchanged proteins more frequently than others (Fig. 3C). Notably, we observed a reorganization of the translational machinery during adipogenesis. In mature adipocytes, the N-terminal acetyltransferase C (NATC) complex was associated with ribosomes, while in preadipocytes, the subunits of the same complex exhibited a diffuse distribution across all fractions (Figure 3D and 3E). Indeed, NATs can bind to ribosomes where they perform N-terminal acetylation in a co-translational manner to regulate protein degradation rates and interactions^12^. Strikingly, among NAT complexes, NATC is particularly important to modify mitochondrial proteins, which are strongly induced in adipogenesis^13^. As another example, we mapped the translocation of numerous mitochondrial proteins, including deoxythymidylate kinase DTYMK, which is involved in pyrimidine biosynthesis. During adipogenesis, DTYMK displayed increased mitochondrial targeting, with a concomitant decrease in the cytosol (Fig.3F).

Additionally, we observed an alternative mechanism contributing to changes in protein expression during adipogenesis, which involved the specific expression of isoforms with distinct localizations. This phenomenon was observed for SLC25A10, the mitochondrial dicarboxylate carrier responsible for succinate transport and predominantly expressed in white adipose tissue^14^. During adipogenesis, isoform 2 with a nuclear profile was downregulated, while isoform 1 with mitochondrial localization significantly increased (Figure 3F, G and H). We further confirmed the isoform switch-driven relocalization through immunofluorescence staining of SLC25A10 using an antibody recognizing both isoforms. The staining showed SLC25A10 localization in the nucleus of preadipocytes and in mitochondria of adipocytes (Figure 3I). In summary, our findings demonstrate that approximately 20% of all mapped proteins undergo localization changes, suggesting that the regulation of protein localization contributes to cellular differentiation processes during adipogenesis.”

### Cooperative protein localization and abundance changes drive metabolic reprogramming

To gain a better understanding of how organelles respond during adipogenesis, we conducted cluster analysis of temporal protein profiles assigned to specific organelles as exemplified here for mitochondria. Our analysis indicated that the notable increase in the total amount of mitochondrial protein during adipogenesis (Fig.3C) was accompanied by the upregulation of various mitochondrial pathways, including the TCA cycle, respiratory chain complexes, and branched chain amino acid (BCAA) catabolism (Fig.4A), consistent with previous findings that degradation of the amino acids valine, leucine, and isoleucine provides an essential pool of acetyl-CoA for de novo lipogenesis in adipocytes^15^. Our spatiotemporal data integration revealed compartment-specific regulation of both the levels and localization of BCAA catabolism enzymes (Fig.4B). Specifically, we found that during the proliferative phase of adipocyte precursor cells, BCAT1 and BCAT2, the first enzymes in the BCAA degradation pathway, are in the cytosol. This localization enables the degradation of BCAA to produce glutamine, a key component required for de novo nucleotide biosynthesis, which is critical for cell division. However, during differentiation, BCAT1 was downregulated, whereas BCAT2 was upregulated and translocated to the mitochondria (Fig.4C). As all mitochondrial BCAA enzymes increase their levels during differentiation, we hypothesize that the upregulation of these enzymes, coupled with BCAT2 translocation to the mitochondria, shifts the pathway from cytosol to mitochondrial BCAA degradation, leading to the production of acetyl-CoA via the TCA cycle, which is essential for de novo lipogenesis in adipogenesis.

**Figure 4.**
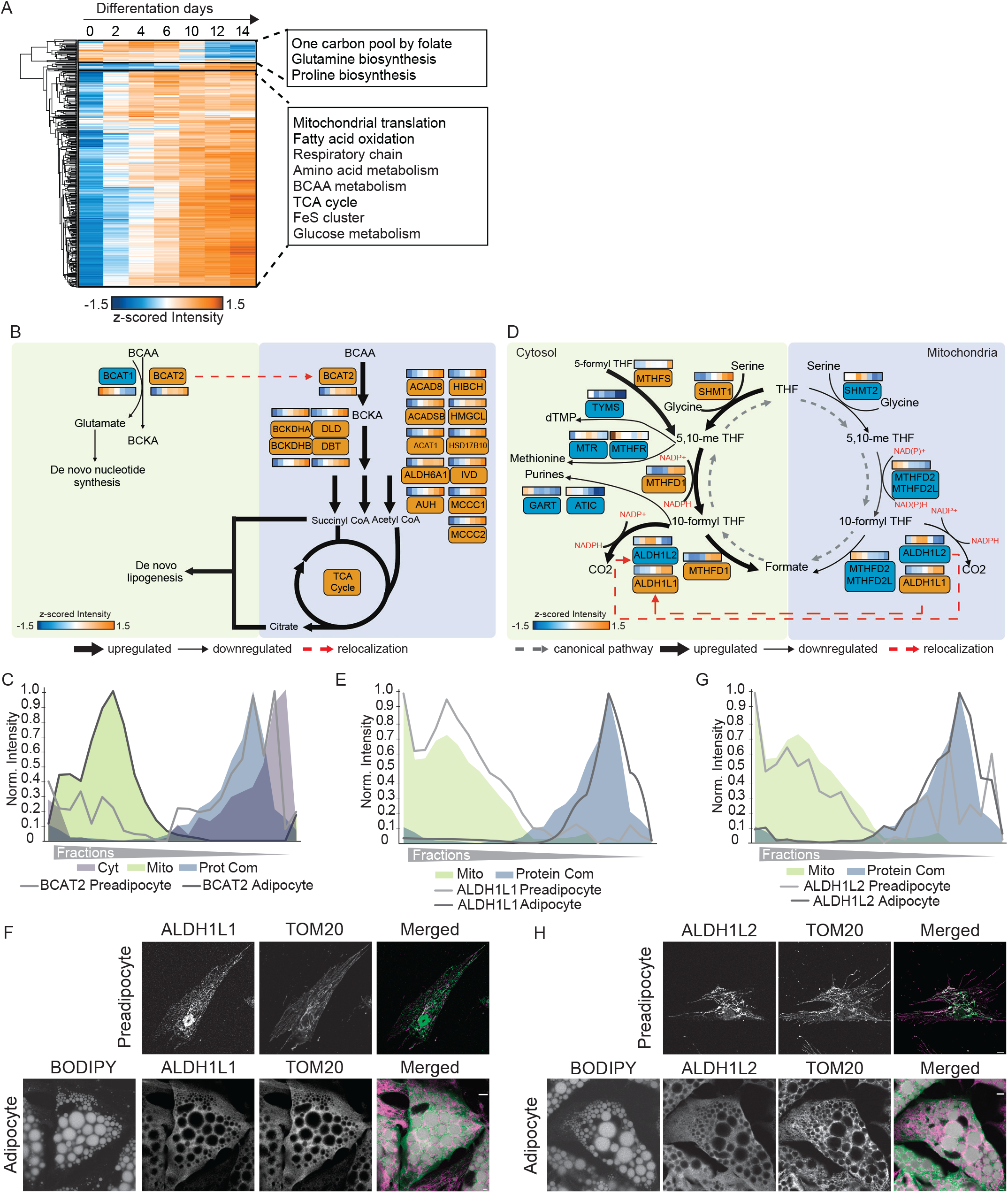
Integration of spatial proteomics with protein levels to characterize organelle metabolic reprogramming. (A) Hierarchical clustering of significantly changed protein profiles in total proteome measurements over the differentiation time course for all proteins predicted as mitochondrial. GO-annotation and Keywords enriched in the clusters are highlighted. (B) Scheme for BCAA metabolism. Up-and down regulated proteins are marked in orange and blue, respectively. Predicted flux and protein localization changes are indicated by arrows. (C) Protein profile of BCAT2 and indicated organelle marker profiles. (D) Scheme of one-carbon metabolism. Up-and down regulated proteins are marked in orange and blue, respectively. Predicted flux and protein localization changes are indicated by arrows. (E) and (G) Protein profiles of ALDH1L1 and ALDH1L2 and indicated organelle marker profiles. (F) and (H) Immunofluorescence for ALDH1L1 and ALDH1L2 in preadipocytes and adipocytes, respectively. In the overlay, BODIPY is shown in grey, ALDH1L1 and ALDH1A2 in green and TOM20 in magenta. Scale bar = 25µm and 10µm in inlay.

Unexpectedly, the increase in mitochondrial protein mass during adipogenesis was accompanied by a decline in the levels of mitochondrial enzymes involved in one-carbon metabolism, which activates and transfers one-carbon units for biosynthetic processes. Given the counter-regulation of this set of enzymes to the mitochondrial protein pool, together with the observation that enzymes for 10-formyltetrahydrofolate catabolism were enriched among the protein with localization changes (Fig.S3B), we investigated the reorganization of one-carbon metabolism during adipogenesis. While levels of all mitochondrial enzymes of the pathway decreased, the cytosolic branch of the pathway was upregulated (Fig.4D). Meanwhile, both isoforms of 10-formyltetrahydrofolate dehydrogenase, ALDH1L1 and ALDH1L2, which catalyze the last reaction of the pathway and promote NADPH release, changed localization from the mitochondria to cytosolic protein complexes, as indicated by their protein profiles, as well as confirmed by co-immunostaining with the mitochondrial marker TOM20 (Fig.4E, F, G and H). Cytosolic enzymes for purine and methionine synthesis catalyzing reactions consuming one carbon metabolism intermediates and cytosolic NADPH decreased. Notably, in proliferating cells, the electrochemical potential difference between mitochondrial NADH and cytosolic NADPH is responsible for driving the serine cycle in the direction that catabolizes serine in the mitochondria and synthesizes it in the cytosol, as shown in previous studies^16^. However, when the mitochondrial component of the serine cycle is lost, the direction of the cytosolic part of the cycle is reversed, leading to cytosolic serine degradation and NADPH^17^ production. Given this regulatory mechanism, compartment-specific adjustments of enzyme levels during adipogenesis may also lead to an increase in cytosolic NADPH synthesis, thereby supporting de novo lipogenesis by providing the necessary reduction equivalents. Compartment-specific regulation of one-carbon metabolism and BCAA enzyme levels was also present in pACs compared to SVF (FigS3C and S3D), indicating that metabolic reprogramming also occurs in vivo during adipogenesis. Together, our findings highlight the coordinated control of protein localization and levels to reprogram metabolic pathways to provide building blocks and reduction equivalents for fatty acid synthesis in adipogenesis.

### Spatial organization of lipid metabolism in human white adipocytes

The defining feature of white adipocytes is their specialization for lipid storage and the formation of large LDs. Therefore, we used our spatial-temporal atlas to investigate the organization of lipid metabolism and to define the adipocyte LD proteome. Hierarchical clustering of the protein profiles had revealed that proteins organized into protein complexes were clearly separated from cytosolic proteins using the PCP approach (Fig.2C). Unexpectedly, annotation enrichment analysis identified not only the partitioning of several prominent complexes, including mTOR, proteasome or chaperonin complexes, into this protein complex cluster, but also an enrichment for fatty acid biosynthesis (Fig.5A). Notably, the enzymes ACLY, FASN, ACACA, and ACACB, which are involved in all steps of fatty acid biosynthesis, from citrate via acetyl-CoA and malonyl-CoA to the final product palmitate, were also sorted into this cluster. Their protein profiles were nearly identical (Fig.5B), suggesting potential condensate formation or a special arrangement of these enzymes within the cytosol in adipocytes.

**Figure 5.**
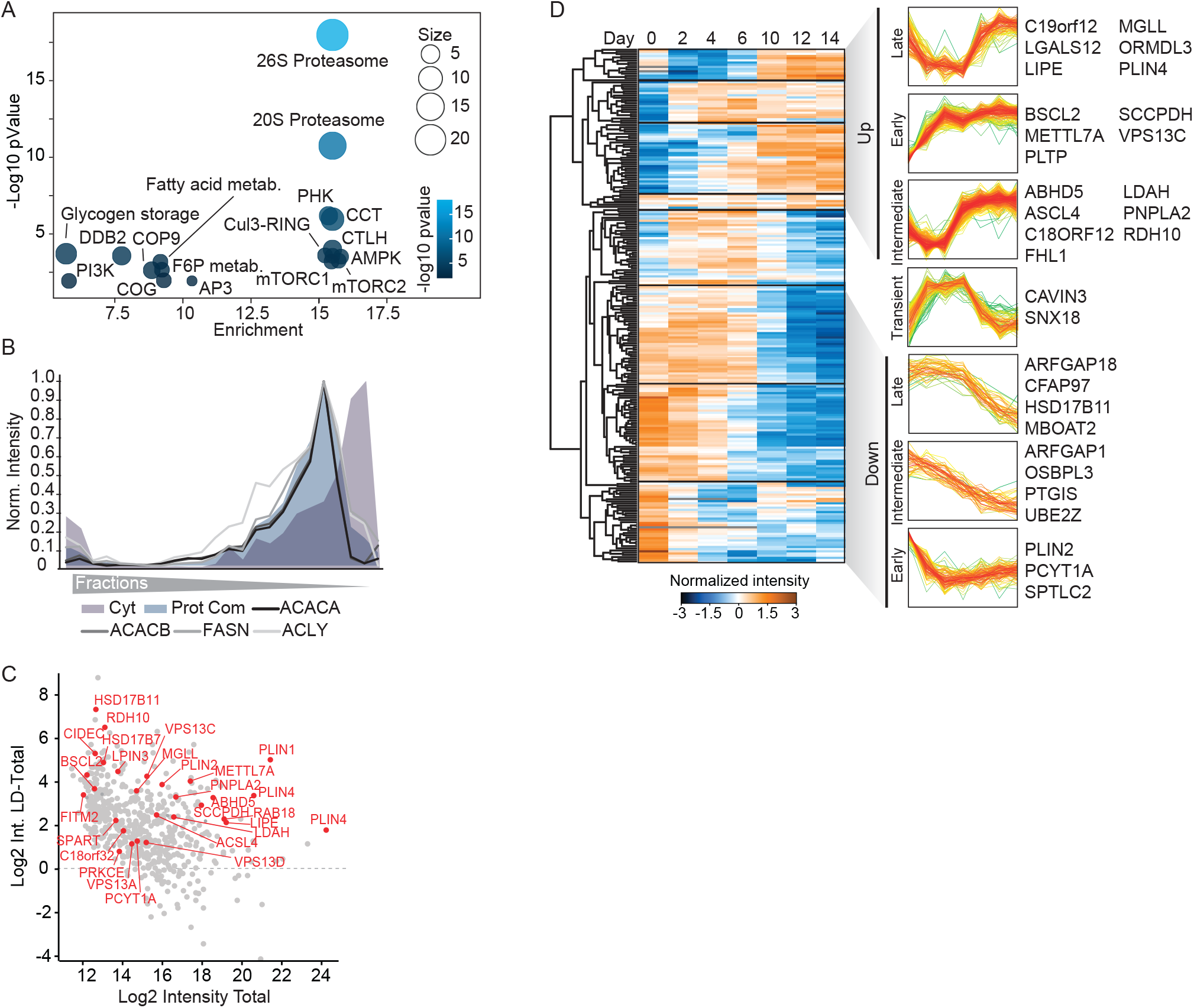
Spatial Organization of lipid metabolism in adipocytes. (A) Corum and Keyword enrichment analysis of proteins identified in the protein complex cluster of the protein correlation profiling in adipocytes. (B) Profiles of indicated enzymes involved in de novo fatty acid synthesis from citrate overlaid with median profile of cytosolic and protein complex associated proteins. (C) Filtering of LD assigned and enriched proteins versus the total proteome. (D) Hierarchical clustering of temporal profiles of protein levels of LD proteins during adipogenesis. Clusters with distinct temporal responses are indicated and examples of proteins found in these clusters are shown. P values and enrichment scores are indicated.

Although adipocytes are the major cell type for lipid storage, a high confident adipocyte LD proteome is lacking so far. To establish the human white adipocyte LD proteome and to exclude contaminants from the set of proteins with LD classifications, we filtered for proteins that were enriched in the LD fraction compared to the total proteome, as all known LD marker proteins showed this behavior (Fig.5C). Clustering analysis of significantly altered LD proteins revealed a well-coordinated and time-dependent regulation of the LD proteome during adipogenesis, which exhibited high consistency across all models examined (Fig.5D). Following adipogenesis induction, a rapid surge in seipin (BSCL2) levels was observed, highlighting its crucial role in the budding of LDs from the endoplasmic reticulum^18–20^. In contrast, the proteins involved in lipid mobilization displayed an upregulation pattern in the later stages. Among the canonical lipolysis-associated proteins, PNPLA2 and its essential cofactor ABHD5 exhibited coordinated upregulation and were assigned to a cluster showing intermediate upregulation. Conversely, LIPE and MGLL were upregulated during the later stages of adipogenesis. Furthermore, alterations in the expression levels of perilipins, a family of structural proteins responsible for coating LDs and regulating lipase access^21^, were also observed. Initially, the ubiquitously expressed member of the perilipin family, PLIN2, was downregulated, whereas the adipocyte-specific member PLIN4 was upregulated late in the adipogenesis process.

### C19orf12 is an adipocyte specific LD protein regulating lipid storage

To identify adipocyte specific LD proteins as potential candidates to promote the exceptional characteristics of adipocytes for lipid storage and dynamics, we overlaid the LD proteome from SGBS cells with the LD proteome from hAPCs, which we also generated by SVM-based protein correlation profiling and subsequent filtering for LD enrichment versus the total proteome (Fig.S4A, S4B). In the adipocyte LD proteomes, we detected most known LD proteins with most of these proteins involved in triglyceride and sterol metabolism (Fig.S4C). We identified 48 LD proteins that were common to both white adipocyte models, out of which 23 were specific to adipocytes (Fig.6A). These adipocyte-specific proteins were distinguished by their lack of enrichment in LDs in any of the datasets integrated into the LD Knowledge Portal^22^, which encompasses proteomic data from non-adipocyte cell lines and the liver.

**Figure 6.**
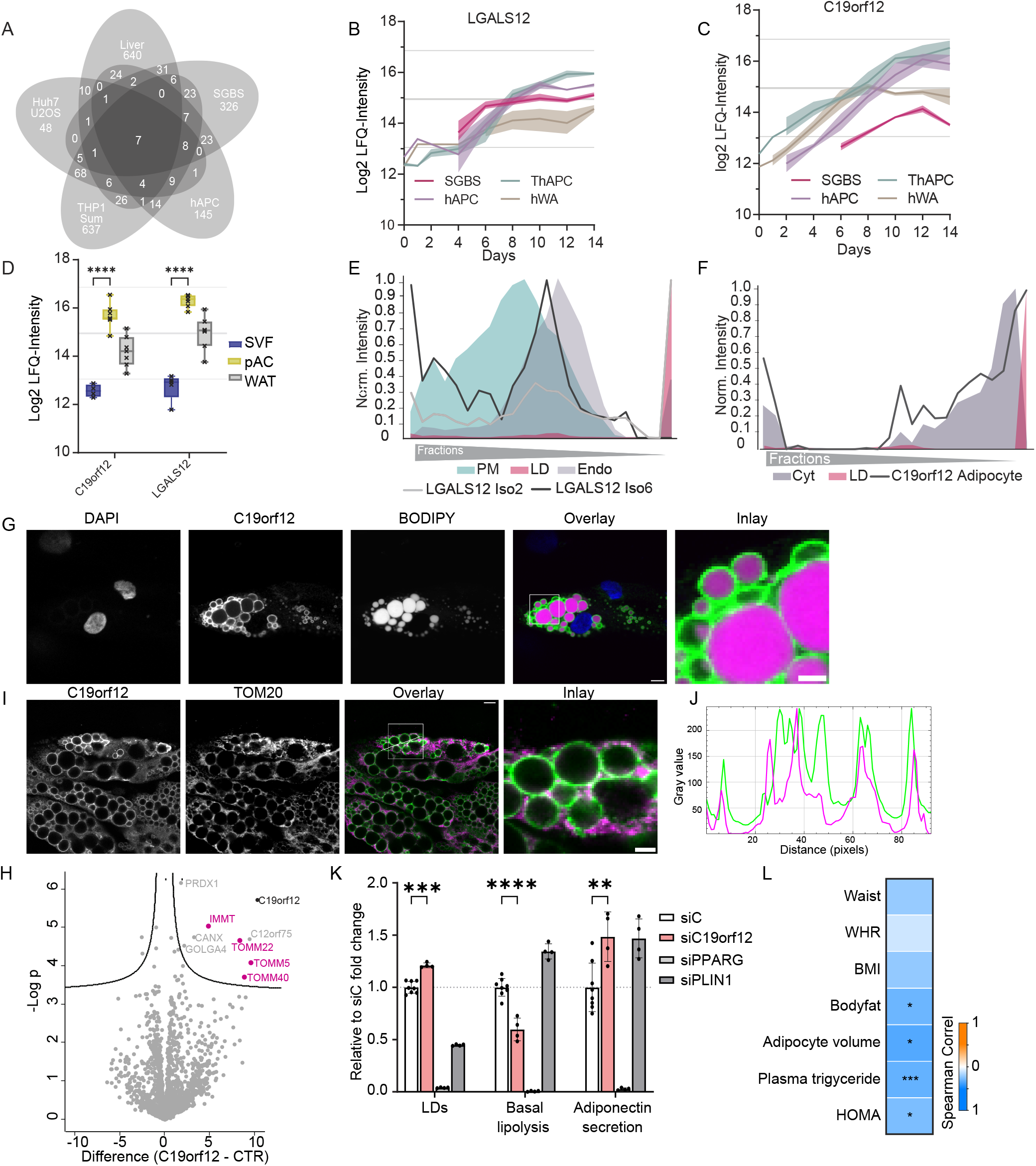
C19orf12 is a LD-mitochondrial contact site protein and regulator of adipocyte function. (A) Venn-diagram showing overlay of proteins enriched in LD fractions versus the total proteome from SGBS and hAPC cells with datasets from the LDKP. (B) Temporal regulation of the level of isoform 2 of LGALS12 during adipogenesis. Isoforms 6 and 7 were not detected in the total adipocyte proteome analysis. (C) Temporal regulation of the level of C19orf12 during adipogenesis. (D) Protein levels in SVFs, pACs and WAT of indicated proteins. Significancy levels represent results from unpaired t-tests with Welch correction and Holm-Šidák adjustment for multiple comparisons, **** = <0.0001. (E) (E) Protein profiles of two detected isoforms of LGALS12 in adipocytes overlaid with LD, plasma membrane and endosomal consensus profiles. Isoforms 6 and 7 display plasma membrane and endosomal localizations, while isoform 2 localizes to LDs. (F) Protein profile of C19orf12 in adipocytes overlaid with indicated organelle marker profiles (G) Immunofluorescence for C19orf12 in SGBS adipocytes. In overlay, BODIPY displayed in magenta, C19orf12 in green and DAPI in blue. Scale bar = 25µm and 10µm in inlay. (H) Volcano plot of the interactome of C19orf12-GFP vs. GFP control in SGBS preadipocytes overexpressing GFP-tagged protein. Components of the mitochondrial protein import machinery are indicated in pink (n=4, FDR<0.05). (I) Immunofluorescence for C19orf12 in SGBS adipocytes. In overlay, TOM20 is displayed in magenta and C19orf12 in green. Scale bar = 25µm and 10µm in inlay. (J) Intensity plot of the fluorescence signal for C19orf21 and TOM20 in the indicated line in (I). TOM20 displayed in magenta and C19orf12 in green. (K) Plot of LD area, adiponectin secretion and basal lipolysis rate for the knockdown of the indicated target genes in hAPC cells on day-1 before differentiation (n= 4-8, error bars represent standard deviation, independent unpaired t-test *p<0.01, ** p<0.001, *** p<0.0001, **** p<0.00001). (L) Association of C19orf12 expression with clinical parameters. Spearman rank correlation test performed on transcriptome analysis of WAT. Color code displays the correlation values (*p<0.01, ** p<0.001, *** p<0.0001, **** p<0.00001).

To delve deeper into their functional significance, we integrated the core proteome data of adipogenesis with the adipocyte-specific LD proteome. Our analysis revealed 10 LD proteins that exhibited significant regulation during adipogenesis. Among these candidates, LGALS12 and C19orf12 particularly captured our attention due to their highly conserved temporal trajectory across all four human adipogenesis models (Fig.6B and C) and their substantial increases in expression exceeded those of other candidate proteins in both cell lines and pACs when compared to the stromal vascular fractions SVFs, suggesting a potential functional role in adipocyte lipid storage (Fig.6D and Fig.S4D). LGALS12, an exclusively adipose tissue-expressed protein, has previously been associated with visceral adipose tissue mass^23^ and demonstrated to influence lipolytic signaling on LDs. Our analysis revealed two isoforms of LGALS12, of which only one isoform (isoform2) being induced during adipogenesis and localizing to LDs (Fig.6E).

C19orf12, is associated with the neurodegenerative disease MPAN (Mitochondrial Membrane Protein-Associated Neurodegeneration)^24^, but has a genetic association with body mass index (Fig.S4E), shows cell type enriched expression for adipocytes^25^ and previous studies indicate a dysregulation in lipid metabolic genes^26^. However, its localization and function in adipocytes remain unknown. To address this question, we conducted a series of experiments to investigate its localization and potential role in lipid storage. Firstly, we confirmed the PCP predicted localization of C19orf12 at LDs (Figure 6F and S4F) by immunofluorescence staining. We observed characteristic ring-like structures formed by C19orf12 around LDs stained by BODIPY (Fig.6G). Next, we performed co-immunoprecipitation coupled with proteomics to map its interactome in both preadipocytes and mature adipocytes. Notably, we identified an interaction between C19orf12 and the TOM complex, which is responsible for mitochondrial protein import (Fig.6H, S4G). This interaction suggests that C19orf12 may function at LDs-mitochondrial interaction sites. Therefore, to examine the subcellular localization of C19orf12 in relation to mitochondria, we conducted co-immunostainings using for C19orf12 and the mitochondrial marker TOM20. These experiments revealed a colocalization of C19orf12 with mitochondria in close proximity to LDs at the contact regions between these organelles (Fig.6I and J).

Next, we sought to explore the functional significance of C19orf12 for adipocyte function and lipid storage at the cellular level. We performed knockdowns by reverse transfection with siRNA in preadipocytes prior to their differentiation, using an optimized protocol^27^. The knockdown resulted in a sustained decrease in C19orf12 protein levels throughout the differentiation process (Fig.S4H). Proteomic characterization of the C19orf12 knockdown cells on day 6 of the differentiation process uncovered a broad dysregulation of lipid metabolism. GO-annotation enrichment analysis revealed an upregulation of mitochondrial and peroxisomal proteins as well as proteins involved in fatty acid synthesis. In addition, proteins associated with lipolytic signaling were enriched among significantly changed proteins. These proteome changes were also reflected in altered lipid dynamics in the mature adipocytes. The quantification of LDs stained with BODIPY revealed a significant ~20% increase in LD accumulation and an approximately 50% higher secretion of adiponectin. Moreover, the knockdown led to a twofold reduction in basal lipolysis (Fig.6L). Remarkably, the cellular phenotype corresponded closely with the data obtained from human patients. Transcriptomic analysis performed on a cohort of 56 female patients^28^ yielded a robust association between *C19orf12* expression and various clinical parameters. Statistical testing revealed a significant inverse correlation between the expression of *C19orf12* and factors such as bodyfat, adipocyte cell volume, plasma triglyceride levels and HOMAR (Homeostatic Model Assessment of Insulin Resistance).

## DISCUSSION

Here, we address a major gap in our current understanding of human adipogenesis, shedding light onto the cellular and metabolic changes that occur during the formation of fat cells. By analyzing temporal proteomics data from multiple models of human adipogenesis and incorporating spatial proteomics, we have uncovered a highly coordinated process of cellular remodeling. Our findings offer a high-resolution view of the sequential changes in protein isoforms, abundance, and organelle organization, shedding light on so far unseen spatial organization of metabolic processes.

One particularly significant finding is the identification of the spatial organization of lipid metabolism enzymes within larger assemblies that exhibit distinct fractionation characteristics compared to soluble cytosolic proteins. Notably, all enzymes involved in de novo lipogenesis display nearly identical profiles in the proteomic co-fractionation, suggesting the condensation of diffusely localized enzymes into discrete foci. Indeed, phase separation is a frequently employed mechanism for controlling the biochemical activity of these enzymes and maintaining metabolic homeostasis. Recent studies have described the structures of human ACACA filaments, revealing that ACACA-citrate dimers polymerize into a polymer with high enzymatic activity in the presence of citrate^29^. However, our work suggests the formation of even larger condensates of all enzymes throughout de novo lipogenesis, which might enable efficient substrate channeling and serve to regulate substrate flux through the pathway.

An important aspect highlighted by our study is the role of protein localization changes in cellular differentiation. We have discovered that a striking 20% of all proteins translocate during adipogenesis, thereby implying that protein translocation may play a more significant and general role in cellular differentiation processes. Several of these events may contribute to cellular specialization for lipid storage. Notably, we have observed the assembly of the NATC complex, responsible for modifying mitochondrial proteins through N-terminal acetylation on ribosomes. Previous research has linked NATC to the modification of mitochondrial proteins, which are highly upregulated during adipogenesis^13^. This suggests that the assembly of the NATC complex may play a supportive role in facilitating adipogenesis by ensuring the proper modification of mitochondrial proteins.

By addressing the previously unexplored spatial aspects of adipogenesis, our study has revealed a sophisticated remodeling of protein localization and abundance in pathways that contribute to de novo lipogenesis. Notably, we observed a diversion of BCAA catabolism away from cytosolic nucleotide biosynthesis towards mitochondrial degradation for citrate and acetyl-CoA synthesis. The translocation of BCAT2 from the cytosol to mitochondria, where it assembles with downstream enzymes into a metabolome complex, may enhance the flux through the mitochondrial part of the pathway^30^, which is highly upregulated during adipogenesis. Similarly, we have identified a counter regulation and localization changes of enzymes involved in mitochondrial and cytosolic one-carbon metabolism. This coordinated regulation may result in the redirection of the cytosolic part of the cycle towards serine degradation and NADPH production, similar as it has been reported for the ablation of mitochondrial one-carbon metabolism enzymes^17^ and thereby contributing to cellular NADPH pools necessary for providing reduction equivalents for fatty acid synthesis

Our spatiotemporal atlas of adipogenesis has revealed novel factors that are crucial for adipocyte biology. We uncovered that C19orf12 localizes specifically to the contact sites between lipid droplets (LDs) and mitochondria, where it interacts with the mitochondrial protein import machinery. These LD-mitochondrial contact sites have been recognized for their importance in facilitating the transfer of fatty acids released from LDs into mitochondria for beta oxidation or reciprocally, providing mitochondrial-generated ATP for triglyceride storage. Moreover, mitochondria associated with LDs have been shown to possess unique functional and proteomic characteristics. Given C19orf12’s simultaneous interaction with LDs and the mitochondrial protein import machinery, it suggests a potential role for C19orf12 in modulating the protein composition of LD-associated mitochondria in adipocytes. Interestingly, our findings diverge from those of other LD-mitochondrial contact site proteins identified in adipocytes. Whereas the knockdown of other LD-mitochondrial contact site proteins resulted in decreased adipogenesis and lipid accumulation^31^, knockdown of C19orf12 enhanced lipid accumulation and reduced lipolysis. These distinct outcomes further imply the existence of diverse types of organelle contacts within adipocytes, each playing a distinct role in either promoting de novo lipogenesis or facilitating lipid mobilization. The inverse correlation observed between *C19orf12* expression and factors related to adiposity and insulin resistance in a human patient cohort further validate the relevance of our in vitro knockdown experiments but also highlight the clinical significance of C19orf12 in human metabolism. Mutations in C19orf12 are associated with the neurodegenerative disease MPAN characterized by brain iron accumulation^24^. The importance of the finding for MPAN will be investigated in follow up studies focusing specifically on patients’ derived samples.

In summary, we have generated a comprehensive cellular map of human adipocytes, encompassing annotations for 8,000 proteins and their changes during adipogenesis. This resource offers researchers in the fields of lipid droplets and adipogenesis a platform to analyze protein expression, metabolic pathways, and organelle composition throughout adipogenesis.

## DATA AVAILABILITY

Source data for each of the figures are provided with this paper. Proteomic raw data and Spectronaut search tables are available via ProteomeXchange under the identifier PXD043338.

## MATERIALS AND METHODS

### Human sample acquisition

Samples from subcutaneous abdominal WAT was obtained by needle aspiration under local anesthesia (as described^32^) from five women and two men (mean± standard deviation for age 60.7±3.5 years and body mass index 28.7±6.6 kg/m^2^). Mature fat cells and SVF were isolated by collagenase digestion as previously described^4^. All studies were approved by the Regional Board of Ethics in Stockholm and all participants provided informed written consent.

### Cell immortalization

To immortalize hAPCs, we stably integrated the EF-1 alpha promoter driving the expression of Puromycin N-acetyltransferase-P2A-TERT in the adeno associated virus serotype 1 (*AAVS1*) safe harbor locus. For this, we electroporated hAPCs with pSpCas9(BB)-2A-GFP (#48138, Addgene) containing the AAVS1-T2 targeting guide and pSH-EFIRES-P-AtAFB2 (#129715, Addgene), which was digested using BglII [#R0144, NEB] and NotI [#R0189, NEB] and thereafter ligated with Puromycin N-acetyltransferase-P2A-TERT. Two selection steps were carried out to obtain edited cells. First, GFP positive cells were selected using BD FACSAria Fusion (BD Biosciences) and expanded. Second, the cells were incubated in the presence of puromycin [2 µg/mL, #A1113803, Thermo Fisher Scientific].

### Cell culture

Human preadipocytes were cultured and differentiated according to their respective published protocols^2,4,5^. Cell culture was performed in a humidified atmosphere with 5% CO_2_ at 37°C. Details about the differentiation cocktails are specified in Fig.S1A. For induction of lipolysis, differentiated cells were treated with 10 µM of isoproterenol or forskolin for 25 min after 1 hour starvation in low Glucose DMEM medium supplemented with 2% fatty acid free BSA.

### RNA isolation, cDNA synthesis and real-time qPCR

Total RNA was purified using the NucleoSpin RNA kit [#740955, Macherey-Nagel] and the concentration and purity of the samples were measured using a NanoDrop 2000 spectrophotometer (Thermo Fisher Scientific). Reverse transcription and mRNA measurements were performed with iScript cDNA synthesis [#1708891, BioRad] and iQ SYBR® Green Supermix [#1708882, BioRad] kits, respectively. Relative mRNA levels were calculated with the comparative Ct-method: 2^ΔCt-target^ ^gene^/2^ΔCt-reference^ ^gene^. The following primers were used: *PLIN1* (Fwd: TGGAGACTGAGGAGAACAAG, Rev: ATGTCACAGCCGAGATGG), *LIPE* (HSL; Fwd: AGCCTTCTGGAACATCACCG, Rev: ATCTCAAAGGCTTCGGGTGG), *CEBPA* (Fwd: AGCCTTGTTTGTACTGTATG, Rev: AAAATGGTGGTTTAGCAGAG), *PPARG* (Fwd: CCCAGAAAGCGATTCCTTCAC, Rev: AGCTGATCCCAAAGTTGGTGG). *18s* (Fwd: TGACTCAACACGGGAAACC, Rev: TCGCTCCACCAACTAAGAAC) was used as a housekeeping gene.

### siRNA knockout

Short interfering oligonucleotides (siRNAs) were introduced by reverse transfection using DharmaFECT Transfection Reagent (T-2003-04, Dharmacon) as previously described^33^. Briefly, siRNA was pre-incubated with DharmaFECT in DMEM for 30 minutes. Proliferating hAPC cells were trypsinized, resuspended in DMEM-low glucose (Gibco, 31885-023) supplemented with 10mM HEPES (Gibco, 15630-056) and 0.5% Penicillin/Streptomycine (Gibco, 15140-122) and added to the transfection mixture with a final siRNA concentration of 100 nM. An siGenome SMARTpool siRNA, comprising a mixture of 4 siRNA provided as a single reagent, was used to target C19orf12 (Dharmacon, M-014731-01), while siRNA targeting PLIN1 (Dharmacon, M-019595-01) and PPARG (Dharmacon, M-003436-02) were used as positive controls for adipocyte function measures. siGENOME non-targeting control siRNA pool 1 (Dharmacon, D-001206-13-05) was used as a control.

### Confocal Microscopy

Cells were plated in glass-bottom 96 well plates [96 CG 1.5, #5242-20, Zell Kontakt] and cultured as described above. At days 0, 8 and 12 of differentiation, cells were fixed in PBS supplemented with 4% paraformaldehyde [#SC281692, Santa Cruz Biotechnology] for ten minutes at room temperature and washed three times with PBS. LDs and nuclei were stained by incubating the cells with PBS supplemented with BODIPY 493/503 [1:2,500, # D3922, Thermo Fisher Scientific] and Hoechst [1:5,000, #ABCAAB228551, VWR], respectively, for ten minutes at room temperature. Subsequently, cells were washed three times with PBS. Images were acquired using CREST V3 confocal system (Crest Optics) mounted on an inverted Nikon Ti2 microscope equipped with a Prime BSIexpress sCMOS camera (pixel size 6.5 μm) from Photometrics. A Nikon 20x/0.75 air objective was used to acquire images.

### Adiponectin assay

For the analysis of adiponectin secretion, cell culture media was exchanged on day 12 of differentiation and collected on day 14. Adiponectin levels were determined by ELISA (R&D systems, DRP300) according to the manufacturer’s protocol.

### Basal lipolysis

For analysis of basal lipolysis, cells were plated in black clear bottom CellBind 96 well plates [#10386612, Fisher Scientific] and cultured as described above. On D10, cell culture media was exchanged to phenol red-free DMEM/F12 [#21041-033, Thermo Scientific]. Media was collected on day 12 and glycerol release into the culture media was quantified to determine basal lipolysis as described previously^33^. Briefly, 20ul of collected media or glycerol standards (#G7793-5ML, Sigma-Aldrich) were added to individual wells in a 96 well clear bottom black plate (M5686-40EA, Sigma-Aldrich). 100ul of a mixture of Free Glycerol Reagent (#F6428-40ML, Sigma-Aldrich) and Amplex Ultrared (10737474, Fisher Scientific) were added and incubated for 15 minutes at room temperature before fluorescence measurement in a Varioskan microplate reader (Thermo Fisher Scientific) at Excitation/Emission 530/590nm. Cells were fixed and glycerol release was normalized to cell number (nuclei count), as assessed via Hoechst staining described below.

### LD area

Cells were plated in black clear bottom CellBind 96 well plates [#10386612, Fisher Scientific] and cultured as described above. On day 12 of differentiation, cells were fixed in PBS supplemented with 4% paraformaldehyde [#SC281692, Santa Cruz Biotechnology] for ten minutes at room temperature and washed three times with PBS. LDs and nuclei were stained by incubating the cells with PBS supplemented with BODIPY 493/503 [1:2,500, # D3922, Thermo Fisher Scientific] and Hoechst [1:5,000, #ABCAAB228551, VWR], respectively, for ten minutes at room temperature. Subsequently, cells were washed three times with PBS. Scanning of nuclei and LD was performed using Cellinsight CX5 (Thermo Fisher Scientific) and quantified by employing object (nuclei) and spot (LD) detection algorithm in HCS Studio software (Thermo Fisher Scientific).

### Co-Immunoprecipitations (Co-IP)

SGBS cells in the undifferentiated state or on day 7 of differentiation were electroporated using the Neon™ Transfection System (Invitrogen) with C-terminally tagged target genes or GFP control. Two days after transfection cells were washed twice with ice-cold PBS and scraped in ice-cold Co-IP buffer (10 mM Tris/Cl pH 7.5, 150 mM NaCl, 0.5 mM EDTA, protein inhibitor cocktail (Roche)) supplemented with 0.5% Nonident P40. Lysates were incubated on ice for at least 30 min and then cleared by centrifugation (4°C, 10min, 17 000 g). The supernatant was then diluted using 3x the volume of Co-IP buffer without detergent and incubated with washed GFP-Trap Magnetic Agarose (Chromotek) for 1h at 4°C on an end-over-end rotator. After this, beads were separated from the lysate on a magnetic rack, washed twice with Co-IP buffer + 0.05% NP40 and three times Co-IP buffer without detergent. Proteins were released from the beads using on-bead digest according to Chromoteks protocol with the following alterations. Beads were first incubated with 50µl elution buffer I (2 M urea, 50 mM Tris/HCl pH 7.5, 20µg/µl Trypsin (Sigma, t6567), 1 mM DTT) and incubated at for 30 min at 37°C, 1300 rpm. After transferring the supernatant to a new cup, beads were incubated again in 50 µl elution buffer II (2 M urea, 50 mM Tris/HCl pH 7.5, 5 mM chloroacetamide (CAA)) in the same conditions but in the dark. The combined supernatants were then digested overnight at 25°C, 1000rpm. The next day peptides were acidified using 1µl trifluoroacetic acid and purified on C18 stage tips as described before^34^.

### Proteomics

Cells were washed with ice-cold PBS, scraped, boiled for 5 min at 95°C 1000rpm in 2% SDC buffer (2% SDC, 100 mM Tris-HCl pH=8.5) and sonicated (Diagenode Bioruptor, 15 * 30s at high intensity). Protein concentration was determined via BCA Protein Assay (Thermo, 23225). Prior digested overnight (37°C, 1000 rpm) with a 1:50 ratio (protein:enzyme) of trypsin (Sigma, t6567) and LysC (Wako, 129-02541), proteins were reduced and alkylated with 10 mM TCEP and 40 mM CAA at 40°C in the dark for 10min. peptides were acidified by adding equal 1:1 (v:v) of isopropanol, 2% TFA. After centrifugation for 10 min at 15,000 rpm, supernatants were loaded onto activated triple layer styrenedivinylbenzene–reversed phase sulfonated STAGE tips (SDB-RPS; 3 M Empore). Peptides were washed with 100 µl ethylacetate 1% TFA, 100 µl 30% Methanol 1% TFA and 150 µl 0.2% TFA and eluted with 60 µl elution buffer (80% ACN, 5% NH_4_OH). Peptides were lyophilized and dissolved in 10 µl MS loading buffer (2% ACN, 0.1% TFA).

### LC-MS/MS analysis

LC-MS/MS analysis 500 ng of peptides was performed on a Orbitrap Exploris 480 (Thermo Fisher Scientific) equipped with a nano-electrospray ion source and FAIMS (CV50) coupled to an EASY-nLC 1200 HPLC (all Thermo Fisher Scientific). Peptides were separated at 60°C on 50 cm columns with an inner diameter of 75 μm packed in-house with ReproSil-Pur C18-AQ 1.9 μm resin (Dr.Maisch GmbH). The peptides were separated over 1h by reversed-phase chromatography using a binary buffer system consisting of buffer A (0.1 formic acid) and buffer B (80% ACN, 0.1% formic acid). Starting with 5% of buffer B, this fraction was increased stepwise to 45% over 45 min followed by a wash-out at 95%, all at a constant flow rate of 300 nl/min. Peptides were ionized and transferred from the LC system into to the gas phase using electrospray ionization (ESI). A data independent (DIA) tandem mass spectrometry 1h method was used. One ms1 scan (300-1650 m/z, max. ion fill time of 45 ms, normalized AGC target = 300%, R= 120.000 at 200 m/z) was followed by 66 ms2 fragment scans of unequally spaced windows (fill time = 22 ms, normalized AGC target = 1000%, normalized HCD collision energy = 30%, R= 15.000).

### PCP

PCP was performed as previously published with minor modification. PCP was performed according to ^69^ with minor modification. For differentiated SGBS and hAPC cells, 5×150 mm dishes of cells at day 20 post differentiation were used per replicate. For preadipocyte gradients, 3x T175 flasks of SGBS cells at one day post confluency were used. Cells were homogenized with a tissue homogenizer on ice in a 1:1 mixture of scraped cells to lysis buffer (20% sucrose, 20 mM Tris pH 7.4, 0.5 mM EDTA, 5 mM KCl, 3 mM MgCl2, protease inhibitor and phosphatase inhibitor cocktail (Roche). 2 ml supernatant was loaded onto the top of a continuous 11 ml 20%-55% sucrose gradient in 20 mM Tris pH 7.4, 0.5 mM EDTA, 5 mM KCl, 3 mM MgCl2. Organelles were separated by sucrose-density centrifugation at 100.000 g (Beckmann, Rotor SW40 Ti), for 3 hrs at 4°C. To isolate LDs, the 1 ml top fraction was cut with a tube-slicer (Beckman coulter). The underlying 0.5 ml gradient fractions were collected from the top to the bottom of the gradient for proteomic analysis.

### Data Analysis

DIA Raw files were demultiplexed with Spectronauts HTRMS converter and analyzed with Spectronaut (v15.7.220308.50606). Proteomic data was analyzed using MaxQant Version Bioinformatics analysis was performed with Perseus, Rstudio, Circos Table Viewer and Microsoft Excel. Annotations were extracted from UniProtKB, Gene Ontology (GO), the Kyoto Encyclopedia of Genes and Genomes (KEGG), CORUM.

### Human genetic association analysis

Human genetic association analysis was performed using MAGMA (Multi-marker Analysis of GenoMic Annotation)^35^ scores in the LDKP^22^. MAGMA scores for genes of proteins with lipid-droplet localization during adipogenesis were plotted for adiposity-related phenotypes as indicated in the figures. A significance threshold of p < 2.5 × 10^−6^ is generally considered significant for MAGMA. A more stringent significance threshold of p < 3.125 × 10^−7^ was derived by Bonferroni correction to account for the 8 phenotypes tested. Manhattan plots were generated in R using ggplot2.

### C19orf12 Clinical Correlations

Expression level of C19orf12 was retrieved from a previously published WAT microarray study compromising 56 women with or without obesity^28^. This microarray data is publicly available in the NCBI Gene Expression Omnibus repository under the accession numbers GSE25401. In brief, C19orf12 expression was correlated to clinical parameters using the cor function in R (method = ”spearman”).

### Proteome Analysis

For proteome analysis quantified proteins were filtered for at least two valid values among three biological replicates in at least one of the conditions. Missing values were imputed from a normal distribution with a downshift of 1.8 and a width of 0.3. Significantly regulated proteins between the three conditions were determined by ANOVA (FDR 0.01) or for two conditions by Student’s T-test (FDR 0.01). Hierarchical clustering, 1D annotation enrichment, and Fisher’s exact test were performed in Perseus. To identify proteins with conserved temporal trajectory, proteins were filtered for temporal profiles whose correlations are positive in all dual comparisons.

### Protein Correlation Profiling Analysis

LFQ intensities for each protein among the organellar fractions were scaled from 0-1. To the fraction with the maximum intensity, the value of 1 was assigned, whereas fractions in which the protein was not quantified were set to 0. For the generation of median protein, the median values from the biological replicates for each fraction were calculated and a second 0-1 scaling step was performed. Pearson correlations between profiles of biological replicates were calculated for each protein and proteins with a correlation <0 in any of the comparisons were discarded.

### SVM-Based Organelle prediction

Marker proteins were selected based on their documented GO-annotations and robust quantification in our dataset for main cellular compartments and protein complexes. The set was used for parameter optimization and training of the SVM based supervised learning approach implemented in Perseus software ^36^. Parameters were set to Sigma=0.2 and C=8. For SGBS mature adipocytes, four and for the preadipocytes three and hAPC cells one biological replicate were used for organelle classifications using the second organelle assignment option in the Perseus software Version 1.5.6.2. Organelle predictions were filtered for positive assignment to at least one organelle. The correlation value between the experimentally determined protein profile and the assigned in silico generated combination profile is given as measure for the quality of organelle assignment. The alpha value (0-1) is a quantitative measure for the second organelle contribution. Proteins with alpha=0 were defined as single organelle localizing proteins.

## ACKNOWLEDGEMENTS

We thank Eva Maria Trautmann and all members of the Krahmer and Mejhert and Rydén labs for discussions and critical reading of the manuscript. We thank Daniel Brandt and Lukas Wolterek for technical assistance. These studies were supported by DFG Emmy Noether (KR5166/2 to N.K.), DFG BATenergy (TRR 333*/1* – 450149205 to N.K.) and the European Foundation for the Study of Diabetes (Future Leader Award NNF20SA0066171 to N.K. and NNF22SA0081233 to N.M.). This work was also supported by grants from the Swedish Research Council (M.R., N.M), ERC-SyG SPHERES (856404 to M.R.), the Novo Nordisk Foundation (including the MeRIAD consortium Grant number 0064142 to M.R. and NNF20OC0061149 to N.M.), Knut and Alice Wallenbergs Foundation (including Wallenberg Clinical Scholar to M.R.), the Center for Innovative Medicine (M.R.), the Swedish Diabetes Foundation (M.R.), the Stockholm County Council (M.R.), the Strategic Research Program in Diabetes at Karolinska Institutet (M.R.). S.F.C. is supported by a Novo Nordisk postdoctoral fellowship run in partnership with Karolinska Institutet. M.O.H. is supported by postdoctoral fellowship from the strategic research program in diabetes at Karolinska Institutet. A.C.M. was supported by grants from NBIA Poland and NBIA Disorders Association (US). A.I. was supported by Hoffnungsbaum e.V. and NBIA Suisse.

## Author Contributions

N.K. and F.K. conceived the project. N.K., N.M., M.R., S.F.C. and F.K. designed experiments. F.K. and M.O.H. performed organelle fractionations. F.K. conducted proteomic analyses. N.K., F.K., A.T., P.K. and S.F.C. analyzed data. S.R. performed proteomic sample preparations. S.F.C., F.K. and M.O.H. performed the siRNA experiments, qPCRs and microscopy. M.W. provided SGBS cell line. T.D.M. contributed discussions. A.C.M. and A.I. contributed to discussions on the role of C19orf12 in LD accumulation and contact site formation. N.K. and F.K. wrote the manuscript.

## FIGURES

**Figure S1.**
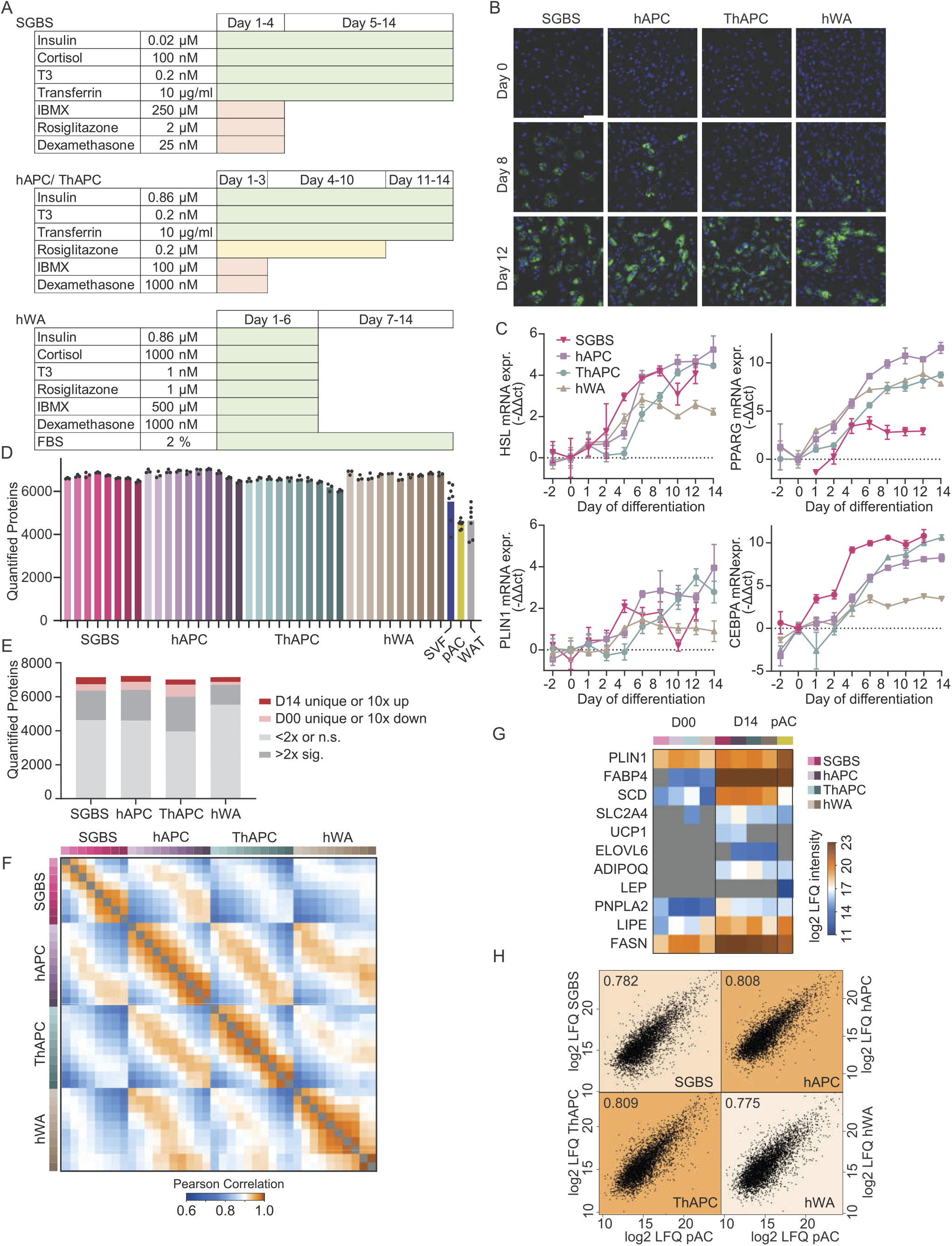
Characterization of human adipogenesis models and overlay with primary cell proteomes. (A) Differentiation cocktails for each cell line. (B) Fluorescent microscopy of cell models at early (Day 0), middle (Day 8) and late stage (D12-D14) of adipogenesis. LDs stained with BODIPY shown in green, nuclei stained with DAPI shown in blue, scale bar 100µm. (C) qPCR of adipogenic marker genes at indicated time points. Values normalized to the first detected time point. (n=4, error bars represent standard deviation). (D) Numbers of quantified protein groups over the differentiation and in primary samples. (Bars represent mean values, dots represent individual replicates). (E) Number of proteins quantified across differentiation in each cell model (min. 2 detections in at least one time-point) and numbers of significantly regulated proteins or proteins exclusively quantified in either mature adipocytes or preadipocytes (Students t-tests, permutation-based FDR<0.05). (F) Pearson correlations of median protein intensities between conditions. (G) Adipose marker protein LFQ intensities at day 0 and day 14 of the cell models and in primary adipocytes. (H) Pearson correlations of median protein intensities between each differentiated model and primary adipocytes (min. 2 valid per condition, missing values imputed with 0).

**Figure S2.**
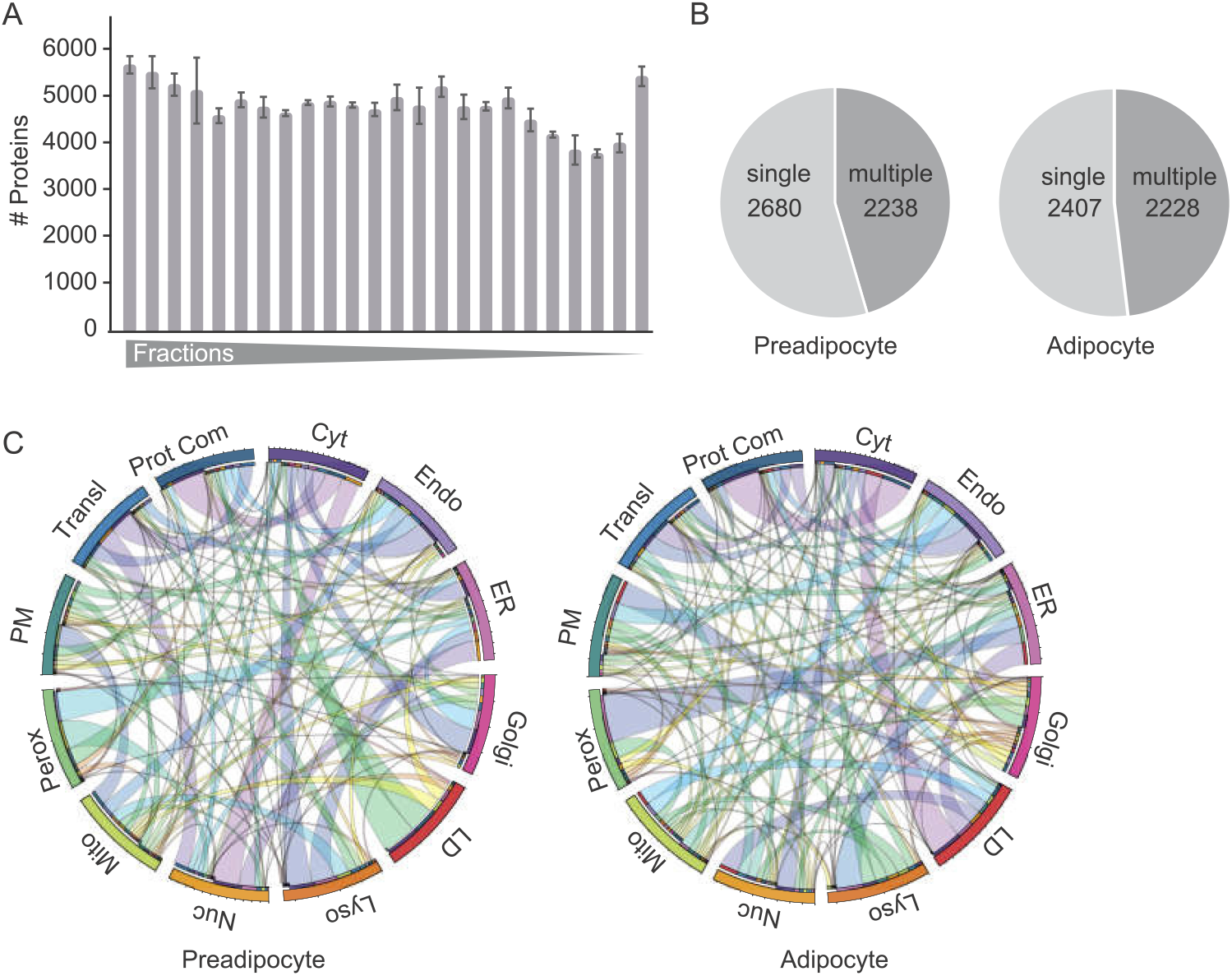
A cellular map of human preadipocytes and adipocytes. (A) Numbers of quantified proteins per organelle fraction in adipocytes (n=4). Error bars represent error of means. (B) Numbers of proteins with single and dual protein predictions in preadipocytes and adipocytes. (C) Circular plot of first and second organelle predictions of preadipocyte and adipocyte organelle maps, respectively. Elevated and resting bars represent first and second assignment, respectively.

**Figure S3.**
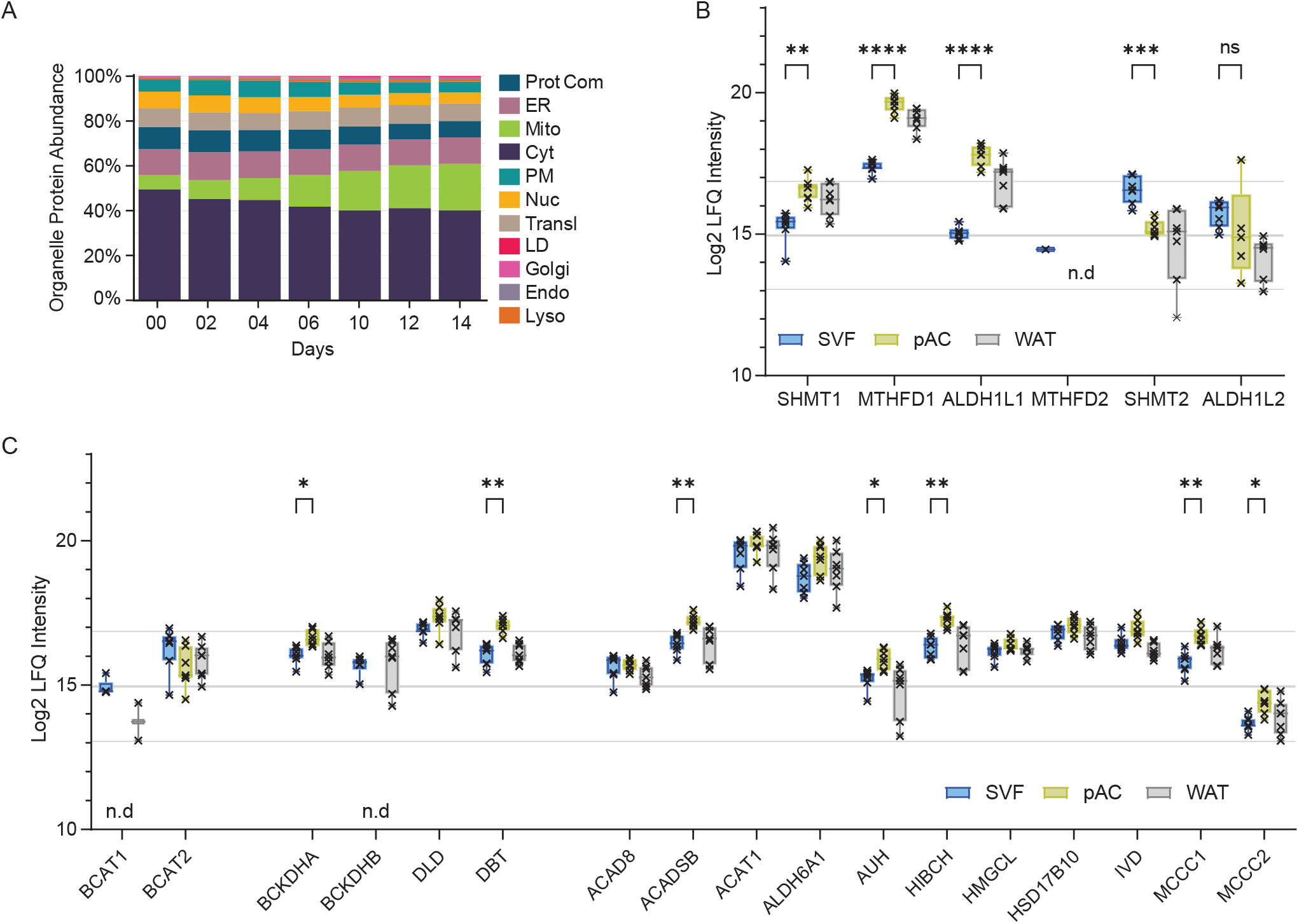
Organelle remodeling during adipogenesis. (A) Percentage of organelle intensities in total proteomes during the time course of differentiation. Calculations are based on first localizations of assigned proteins in either pre- or adipocyte PCP for the respective condition and agreeing localizations for the conditions in between. (B), (C) Protein levels of indicated proteins of BCAA and one carbon metabolism in SVF, pAC and WAT. (Error bars stretch from min. to max. levels, all points shown, for each plot welch corrected unpaired t-tests between SVM and pAC were performed using the Holm-Šidák adjustment for multiple comparisons, * p value<0.05, ** p value <0.01, *** p value <0.001, **** p value <0.0001).

**Figure S4.**
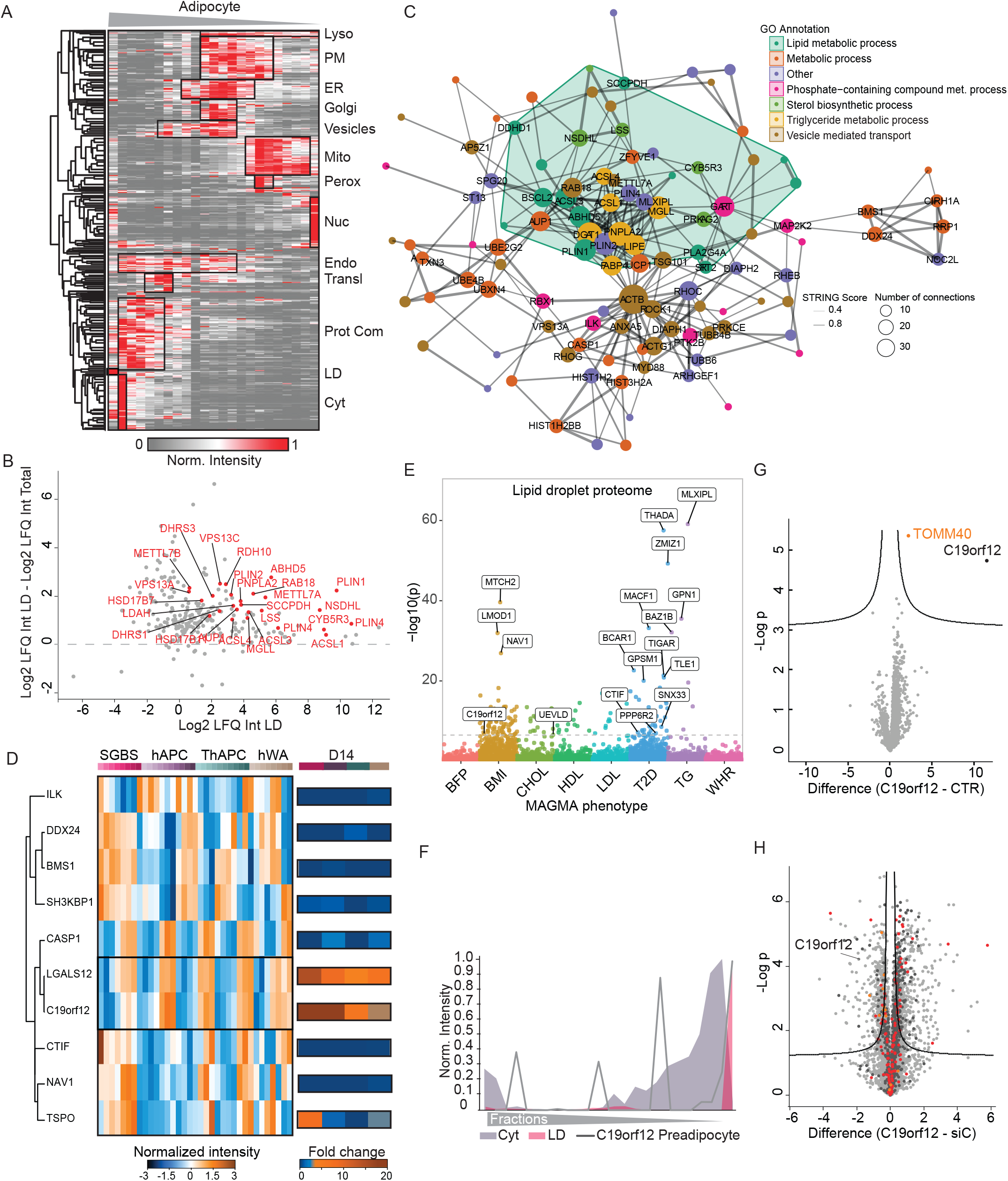
The LD proteome of white adipocytes. (A) Hierarchical clustering of PCP of hAPCs.(n=1). (B) Plot of protein levels from proteins determined as LD proteins as first or second assignments from SVMs based on PCP analysis in hAPCs in LD fraction vs total proteome from hAPCs. (C) LD proteins were mapped in the STRING database, and the ones with at least two partners (physical and/or functional) are shown. High-confidence interactions have thicker connected lines. GO enriched annotation terms (FDR < 0.05) were colored to highlight noteworthy clusters. (Size = number of connections, line width = STRING combined score). (D) Hierarchical clustering of temporal profiles of significantly changed adipocyte specific LD proteins during adipogenesis and their fold change at day 14 of differentiation in four adipogenesis models. (E) LD proteome contains hits associated with metabolic phenotypes. Metabolic associations (−log10 p values) of genes that in SGBS LD proteome and at least one other LD proteome were analyzed for 8 metabolic phenotypes. A significancy threshold value of p = 2.5e-6 was used (dotted line) (F) Protein profile of C19orf12 in preadipocytes overlaid with indicated organelle marker profile. (G) Volcano plot of C19orf12 interactome in differentiated SGBS cells after imputation of missing values (C19orf12-GFP vs. GFP control, n=6, FDR<0.05). (H) Volcano plot of the knockdown proteome of C19orf12 versus siRNA control at D12 of differentiation (n=3, FDR<0.01). Proteins involved in fatty acid metabolism (red), mitochondrial functions (dark grey) and negative regulation of adenylate cyclase activity by GPCR signaling (orange) are indicated.

